# A Complete Public Domain Family Genomics Dataset

**DOI:** 10.1101/000216

**Authors:** Manuel Corpas, Mike Cariaso, Alain Coletta, David Weiss, Andrew Harrison, Federico Moran, Huanming Yang

## Abstract

**Background:** The availability of open access genomic data is essential for the personal genomics field. Public genomic data allow comparative analyses, testing of new tools and genotype-phenotype association studies. Personal genomics data of unrelated individuals are available in the public domain, notably the Personal Genome Project; however, to date genomics family data and metadata are severely lacking, mainly due to cost, privacy concerns or restricted access to Next Generation Sequencing (NGS) technology. Family data have a lot to offer as they allow the study of heritability, something which is impossible to do just by using unrelated individuals.

**Findings:** A whole family from Southern Spain decided to genotype, sequence and analyse their personal genomes making them publicly available under a Creative Commons 0 license (CC0; commonly denominated as public domain). These data include a) five 23andMe SNP chip genotype bed files, b) four raw exomes with their assorted bam files and VCF files, c) a metagenomic raw sequencing data file and d) derived data of likely phenotypes using SNPedia-derived tools.

**Conclusions:** To our knowledge this is the first CC0 released set of genomic, phenotypic and metagenomic data for a whole family. This dataset is also unique in that it was obtained through direct-to-consumer genetic tests. Hence any ordinary citizen with enough budget and samples should be able to reproduce this experiment. We envisage this dataset to be a useful resource for a variety of applications in the personal genomics field as a) negative control data for trait association discovery, b) testing data for development of new software and c) sample data for heritability studies. We encourage prospective users to share with us derived results so that they can be added to our existing collection.

## Data Description

As the price of genome sequencing and all other assorted sequencing techniques become more affordable, this technology is expected to be adopted among the general population in the future. Next Generation Sequencing (NGS) experiments are data rich, but the interpretation of personal genomes remains a challenge as knowledge is spread across many different resources, many of them closed source or with no metadata with which to perform automatic computational analyses. This lack of open source software and tools is in part due to absence of data models with which to test and develop new applications. Family models readily available for development, testing and controls are particularly necessary for the personal genomics field as they are more useful than those of individual genomes, providing the context that affords the research of trait heritability [1].

In this data article, we provide the complete genomics dataset of five blood-related family members that performed a series of direct-to-consumer (DTC) personal genomics tests. To our knowledge this is the first dataset released to the public for a whole family using only DTC means. The genotype data were obtained from 23andMe while all the sequencing data from the BGI. The dataset is summarised in *Table 1*. We call members of this family Mother, Father, Aunt, Daughter and Son. The data presented here include both raw and processed data. The rationale for including raw data, even though most users will not require them, is that current processing tools (e.g. genome aligners; variant call software) may provide different results depending on the type of configuration used. Whenever a type of data can be deduced from another, we have left the former out; e.g. bam indexes can be deduced from bam files so we have not included bam indexes. Also in the case of bam files, files belonging to the same sample have been merged into one. Due to these data having been obtained using DTC means, sometimes the source or platform utilised for the same test may be slightly different between individuals because of a different version of the platform used (indicated in Table 1). We have minimised the impact of platform versions by using the same pipelines and methods for alignment and variant calling.

**Table 1:** Summary of the data presented in this article, belonging to a family of 5 individuals from Southern Europe. The dataset includes a total of 27 files and 18,463 GB of data.

| Type | Provider | Source | Platform | Format | Year Obtained | # Files | Individual(s) | Approx. Size (MB) |
| --- | --- | --- | --- | --- | --- | --- | --- | --- |
| Genotype | 23andMe | Saliva | SNP chip v2 | bed.zip | 2009 | 1 | Son | 5 |
| Genotype | 23andMe | Saliva | SNP chip v3 | bed.zip | 2011 | 1 | Mother | 8 |
| Genotype | 23andMe | Saliva | SNP chip v3 | bed.zip | 2011 | 1 | Father | 8 |
| Genotype | 23andMe | Saliva | SNP chip v3 | bed.zip | 2011 | 1 | Daughter | 8 |
| Genotype | 23andMe | Saliva | SNP chip v3 | bed.zip | 2011 | 1 | Aunt | 8 |
| Annotation | SNPedia | SNP chip v2 | Phenotypes | txt | 2012 | 1 | Son | 0.3 |
| Annotation | SNPedia | SNP chip v3 | Phenotypes | txt | 2012 | 1 | Mother | 0.4 |
| Annotation | SNPedia | SNP chip v3 | Phenotypes | txt | 2012 | 1 | Father | 0.4 |
| Annotation | SNPedia | SNP chip v3 | Phenotypes | txt | 2012 | 1 | Daughter | 0.4 |
| Annotation | SNPedia | SNP chip v3 | Phenotypes | txt | 2012 | 1 | Aunt | 0.4 |
| Whole Exome Sequencing | BGI | Saliva | SE Illumina HiSeq 2000 | fastq.gz | 2011 | 4 | Son | 2400 |
| Whole Exome Sequencing | BGI | Saliva | PE Illumina HiSeq 2000 | fastq.gz | 2013 | 2 | Mother | 2000 |
| Whole Exome Sequencing | BGI | Saliva | PE Illumina HiSeq 2000 | fastq.gz | 2013 | 2 | Father | 2000 |
| Whole Exome Sequencing | BGI | Saliva | PE Illumina HiSeq 2000 | fastq.gz | 2013 | 2 | Sister | 2400 |
| Alignment | InSilicoDB | SE Illumina HiSeq 2000 | BWA | bam | 2013 | 1 | Son | 2900 |
| Alignment | InSilicoDB | PE Illumina HiSeq 2000 | BWA | bam | 2013 | 1 | Mother | 1800 |
| Alignment | InSilicoDB | PE Illumina HiSeq 2000 | BWA | bam | 2013 | 1 | Father | 1600 |
| Alignment | InSilicoDB | PE Illumina HiSeq 2000 | BWA | bam | 2013 | 1 | Daughter | 2200 |
| Annotation | InSilicoDB | Bam from Son, Mother, Father, Daughter | GATK | VCF | 2013 | 1 | Son, Mother, Father, Daughter | 124.6 |
| Metagenomics | BGI | Fecal frozen sample | PE Illumina HiSeq 2000 | fastq.gz | 2013 | 2 | Son | 1000 |
| **TOTAL** |  |  |  |  |  | **27** |  | **18463.5** |

Data available encompass genotypes, annotations, alignments, whole exome sequencing and metagenomics sequencing. Processed data files containing predicted phenotypes, bam files and VCF files [2] are based on the raw data provided (SNP chip, Illumina HiSeq). Although data describing observed phenotypic traits are not included at present, our intention is to provide them in the near future. All variations are SNP-based and no Copy Number Variation (CNV) annotations are provided at this point. Whenever possible, data have been kept compressed to economise data storage. Years in which the data were obtained have also been included to roughly show the timeline of family tests. The five 23andMe SNP chip genotypes are in bed format, mapped to the human reference NCBI36. These datasets were taken in the summer of 2009 (Son; version 2 of 23andMe; ∼0.5M SNPs) and January 2011 (Mother, Father, Daughter, Aunt; 23andMe SNP chip version 3, ∼1M SNPs per individual). Whole exome sequencing from Son, Mother, Father, Sister was done from saliva samples using Illumina HiSeq 2000 with an expected coverage of ∼30x. The Son’s exome was carried out first (January 2012), using the Agilent 44M Single End (SE) capture chip. The other three exomes (Mother, Father, Daughter) were sequenced in January 2013 using the Agilent 44M Paired End (PE) capture chip each. The metagenomics sequencing has been performed for Son in 2013 from a fecal sample using Illumina shotgun sequencing. The processing of phenotypes was performed using Promethease, a tool that takes SNPedia [3] genotype-phenotype associations. Promethease requires 23andMe bed files as input and outputs text files with SNP ids and their annotation. Processing of exome data was carried out with InSilicoDB [4] a web-based storage hub. For all of the exomes, InSilicoDB produced bam files with BWA [5]. Once bam files were created, they were merged into one per individual. InSilicoDB generated variant calls using the Genome Analysis Toolkit (GATK) ‘best practice variant detection method’ pipeline [6]. Variant discovery was performed using joint variant calling, outputting one VCF file for all four processed exomes. All pipelines and data analyses in InSilicoDB are available at https://insilicodb.org/app/browse/?q=ISDB11122. All raw data are available in figShare at http://figshare.com/articles/Corpasome/693052 from which they can be cited.

## Availability and requirements

*Project name*: Corpasome

*Project home page:* http://figshare.com/articles/Corpasome/693052

*Operating system:* Platform independent

*License:* Creative Commons 0

*Any restrictions to use by non-academics*: None

## Availability of supporting data

Processed data are available at https://insilicodb.org/app/browse/?q=ISDB11122. Raw data are available at http://figshare.com/articles/Corpasome/693052.

## Acknowledgements

We are grateful to Fiona Nielsen for useful comments on the manuscript.

